# Dynamic evolution of bacterial ligand recognition by formyl peptide receptors

**DOI:** 10.1101/2021.08.02.454796

**Authors:** Nicole M. Paterson, Hussein Al-Zubieri, Joseph Ragona, Juan Tirado, Brian V. Geisbrecht, Matthew F. Barber

**Affiliations:** Institute of Ecology and Evolution, University of Oregon, Eugene, OR, USA; Department of Biochemistry and Molecular Biophysics, Kansas State University, Manhattan, KS, USA; Department of Biology, University of Oregon, Eugene, OR, USA

## Abstract

The detection of invasive pathogens is critical for host immune defense. Cell surface receptors play a key role in the recognition of diverse microbe-associated molecules, triggering leukocyte recruitment, phagocytosis, release of antimicrobial factors, and cytokine production. The intense selective forces acting on innate immune receptor genes has led to their rapid diversification across plant and animal species. However, the impacts of this genetic variation on immune functions are often unclear. Formyl peptide receptors (FPRs) are a family of animal G-protein coupled receptors which are activated in response to a variety of ligands including formylated bacterial peptides, microbial virulence factors, and host-derived peptides. Here we investigate patterns of gene loss, sequence diversity, and ligand recognition among primate and carnivore FPRs. We observe that FPR1, which plays a critical role in innate immune defense in humans, has been lost in New World primates. Patterns of amino acid variation in FPR1 and FPR2 suggest a history of repeated positive selection acting on extracellular domains involved in ligand binding. To assess the consequences of FPR variation on bacterial ligand recognition, we measured interactions between primate FPRs and the FPR agonist *Staphylococcus aureus* enterotoxin B, as well as *S. aureus* FLIPr-like which functions as an FPR inhibitor. We find that comparatively few sequence differences between great ape FPRs are sufficient to modulate recognition of *S. aureus* ligands, further demonstrating how genetic variation can act to tune FPR activation in response to diverse microbial binding partners. Together this study reveals how rapid evolution of host immune receptors shapes the detection of diverse microbial molecules.

## INTRODUCTION

The formyl peptide receptors (FPRs) are a family of G-protein coupled receptors (GPCRs) that play crucial roles in the recruitment and activation of leukocytes in vertebrates. Early studies demonstrated that certain human cell types migrate towards N-formylated peptides, which are present in bacterial and mitochondrial, but not eukaryotic, proteins (Schiffmann 1975). These findings led to the discovery of FPRs as a new class of pattern recognition receptor with the ability to discriminate between ‘self’ and ‘non-self’ molecules in order to activate downstream immune responses (Pike 1980.; Zigmond 1981). Since then, additional microbial and host-derived ligands have been identified for specific FPR homologs. Of the three FPRs in humans, each has been shown to possess a unique ligand-binding profile (Karlsson 2009.; Schepetkin 2014; Kretschmer 2010), with a range of output responses. For example, recognition of lipoxin-A by FPR2 leads to the suppression of inflammatory signaling, whereas binding of bacterial-specific formylated peptides by FPR1 results in induction of the inflammatory response and cell chemotaxis towards the ligand source (John 2007; Le 2002; Schepetkin 2014).

Human neutrophils and other myeloid cells play a central role in innate pathogen recognition and express high levels of FPRs. When neutrophils detect foreign molecules via FPR activation, the cell migrates towards the source of the signal. Upon reaching a site of infection, neutrophils contribute to pathogen clearance through phagocytosis, release of toxic granule molecules, and a rapid oxidative burst which produces high levels of antimicrobial reactive oxygen species and reactive nitrogen species (Shimizu 2008.; Önnheim 2014; Bufe 2015). Neutrophils constitute roughly 50% of circulating bloodstream leukocytes and are capable of detecting nanomolar concentrations of pathogen-derived peptides through activation of FPR1 and FPR2 (Fu 2006; Le 2002). Natural killer cells, monocytes, and macrophages also express high levels of formyl peptide receptors which similarly contribute to activation and chemotaxis of these cell types (Leslie 2021; Kim 2009; Crouser 2009).

Immune receptors are under constant selective pressure to detect an array of rapidly evolving microbial ligands (Daugherty and Malik 2012, Aleru and Barber 2020). Beneficial mutations that enhance immune responses and improve fitness are expected to spread rapidly in host populations via positive selection. Previous studies have detected signals of positive selection in FPR1 and FPR2 in the mammalian lineage through observation of elevated rates of non-synonymous to synonymous substitutions (dN/dS) at several codon sites (Muto 2015). This is consistent with a model of formyl peptide receptors undergoing positive selection at sites important for immune function, most likely as it pertains to pathogen detection. All three FPR genes are located on chromosome 19 in humans (Bao 1992). Orthologous FPR gene regions in primates have previously demonstrated evidence of accelerated evolution in promoters and other regions (Yang 2014) as well as heightened occurrences of single nucleotide polymorphisms (Harris 2020). Many other immune system related proteins including Toll-like receptors (TLRs), TRIM proteins, major histocompatibility complex (MHC) family genes, and transferrins have been subject to repeated positive selection during mammalian evolution (Brunette 2012.; McLaughlin and Malik 2017; Daugherty and Malik 2012; Barber and Elde 2014; Aleru and Barber 2020; Paterson 2021; Sawyer 2005.; van der Lee 2017.; Hughes and Piontkivska 2008; Hughes and Nei 1990). The central role of FPRs in innate immunity coupled with previous evidence of positive selection suggest there may be important functional consequences for sequence-level variation observed in this receptor family among mammals.

## RESULTS

### Loss of FPR1 in New World primates

To investigate potential consequences of FPR evolution, we first compiled a dataset of FPR homologs from various primate and carnivore species. Unexpectedly, we found evidence for a complete loss of FPR1 expression in New World primates (Yang and Shi) demonstrated by a lack of FPR1 cDNA detectable in whole blood, brain, lung and other RNA-Seq data from AceView (Thierry-Mieg and Thierry-Mieg) (Figure 1A). This loss of expression suggested to us that the *FPR1* gene may have been lost or pseudogenized in New World monkeys. We identified several pseudogenes in New World monkeys that displayed the absence of gene expression described, as well as a lack of annotated or homologous *FPR1* New World monkey genes in the NCBI database and incomplete gene sequence that nonetheless shared sequence identity with related *FPR1* genes (Figure 2). To determine whether the *FPR1* genes in New World monkeys were likely to be pseudogenes or unexpressed or degraded mRNA, we scanned available New World monkey genomes using a BLAT search and found additional regions of similarity to *FPR1* genes. We tested for the presence of predicted exons using MIT’s GENESCAN tool (Burge and Karlin 1997), but failed to identify exons for any of the gene regions at the given probability cut-off for loci in New World monkeys containing significant similarity to marmoset *FPR1*. We aligned regions identified as putative pseudogenes in New World monkeys to the human *FPR1* reference and the marmoset *FPR1* pseudogene sequences (which are available as high coverage, well-annotated entries in the NCBI database). *Sapajus apella*, *Cebus imitator*, *Saimiri boliviensis* and *Aotus nancymaae* have substantial similarity to FPR1 at similar loci in the genome (adjacent to the *FPR2* and *FPR3* genes on chromosome 19) but lack one or more features of a functional gene (Figure 2A, Figure 2B). The *Saimiri boliviensis* pseudogene is the most striking, as this region has the least conservation (77.5% identity to marmoset *FPR1* in 271bp region located on the plus-strand adjacent to the *FPR2* and *FPR3* genes on chromosome 19), lacks evidence of a start or stop codon, and lacks significant sequence identity to marmoset *FPR1* outside of the 271bp region. A caveat to this analysis is that each of these genomes has different levels of coverage and/or one or more builds (details on the quality of genomes used in this study can be found in the Supplemental Figure 2). However, *Saimiri boliviensis*, *Aotus nancymaae* and *Callithrix jacchus* have multiple builds with high coverage (*Saimiri boliviensis*: 2 builds most recent 111x coverage, *Aotus nancymaae*: 4 builds, most recent 132x, *Callithrix jacchus*: 11 builds, most recent 40x coverage) (Supplemental Figure 2). Collectively these findings provide strong evidence for the loss of the major immune receptor FPR1 in New World primates.

**Figure 1.**
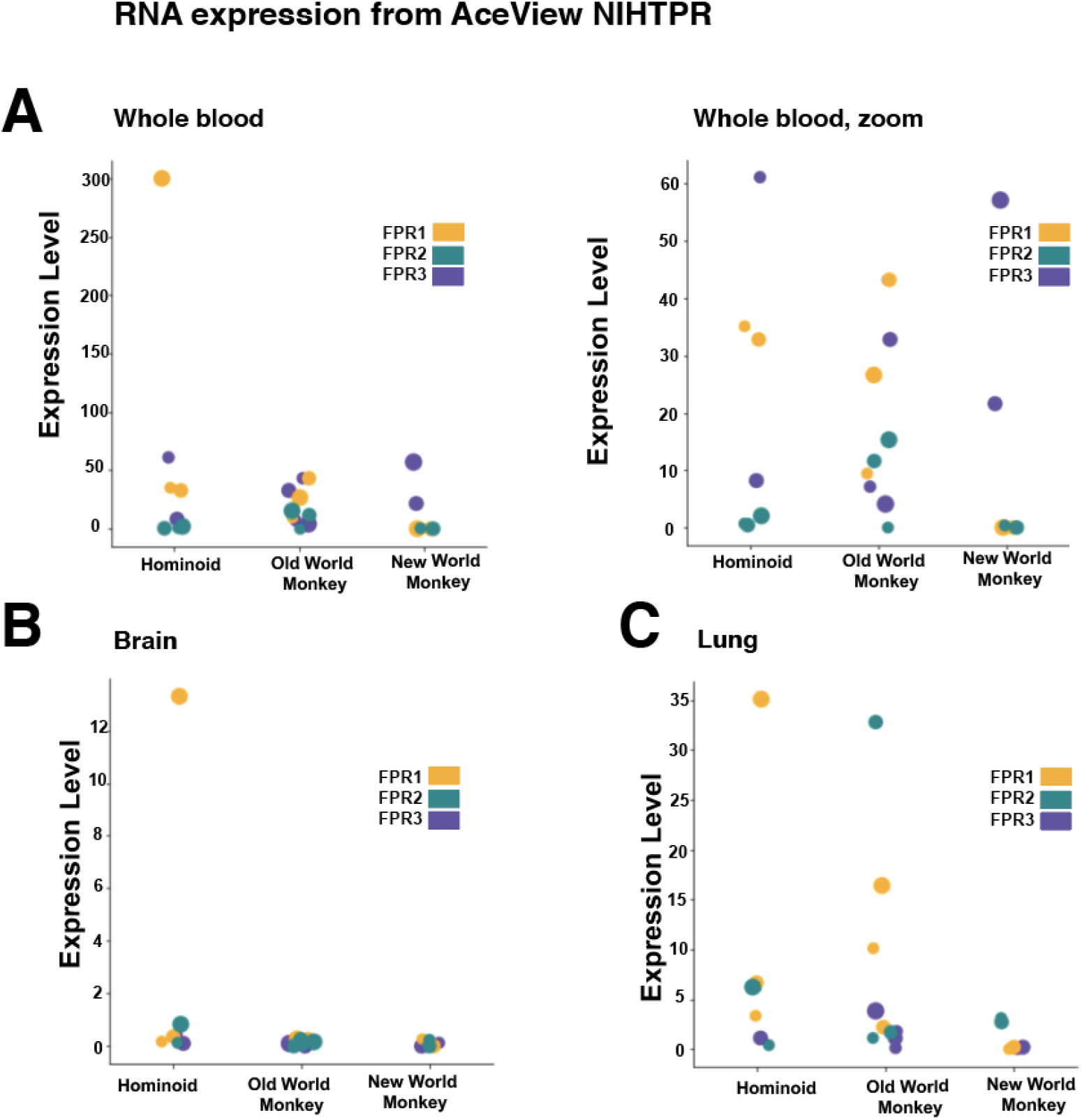
FPR gene expression among simian primates. Aceview NIHTPR RNA-seq data plotted to display FPR expression levels in **A)** whole blood, **B)** brain, and **C)** lungs across hominoid (human, chimpanzee), Old World monkey (pig-tailed macaque, crab-eating macaque, baboon, mangabey) and New World monkey (marmoset, squirrel monkey and owl monkey) species.

**Figure 2.**
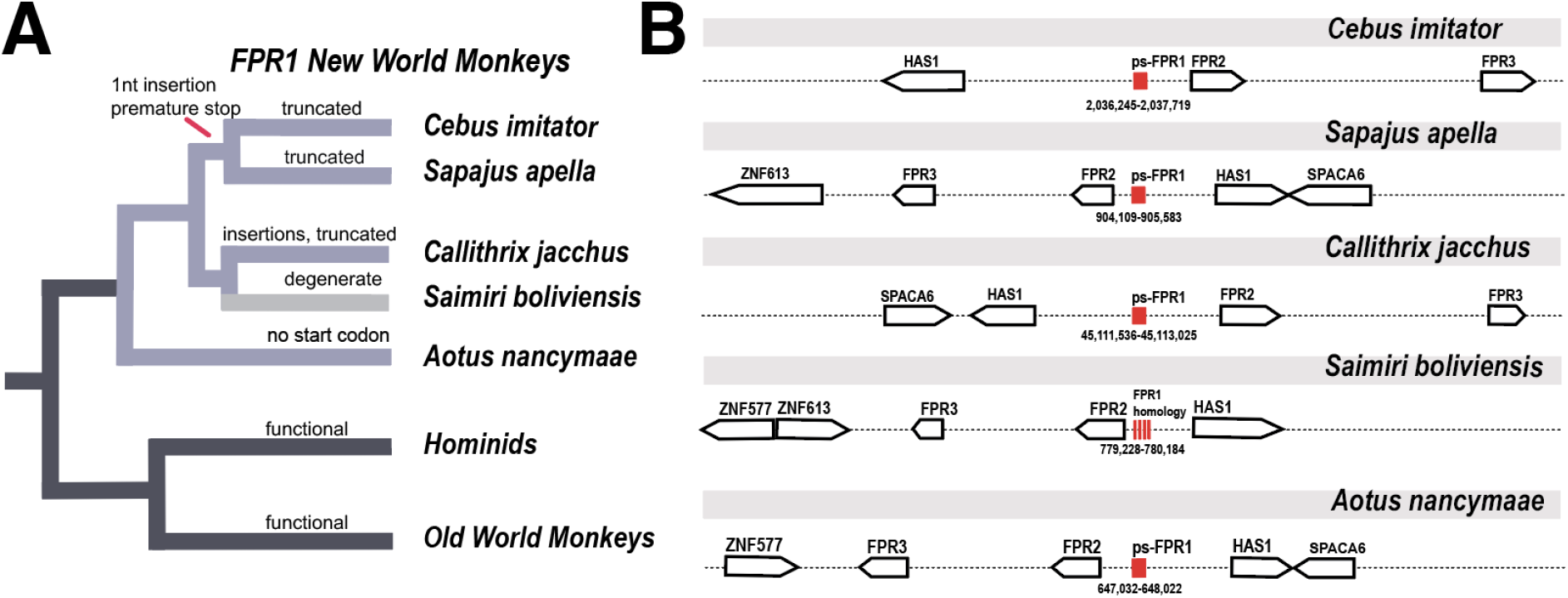
Loss of FPR1 in New World primates. **A)** Evidence for inactivating mutations in FPR1 within the New World monkey lineage. *Saimiri boliviensis* displayed the most significant degeneration in this region, encoding only a short 271 nucleotide sequence with significant FPR1 similarity and no apparent start or stop codons. **B)** Conserved synteny of the FPR gene cluster across New World monkeys.

### Rapid divergence of FPR ligand binding domains in primates and carnivores

The gene expression profile of FPRs in New World monkeys shows substantial expression of FPR2 and decreased expression of FPR1 and FPR3. Recent studies suggest that gene loss in mammals can be an adaptive response to environmental pressures (Monroe 2021; Helsen 2020; Sharma 2018). To test whether sequence divergence in these genes may be the product of natural selection, we first implemented phylogenetic analysis by maximum likelihood (PAML) on primate and carnivore FPR genes (Yang 2007). We included carnivores in addition to primates because carnivores encode only a single FPR homolog, FPR2, and we were curious whether this reduced gene copy number may alter selective forces on this gene compared to other taxa. Our analysis revealed evidence for positive selection acting on FPR1 and FPR2 genes in primates, largely consistent with previous studies across mammals showing heightened dN/dS in FPR1 and FPR2 (Muto 2015) (Figure 3). We performed additional testing for branch-site episodic positive selection using DataMonkey’s abSREL test (Smith et al.) and found evidence of heightened selection on the branch of the phylogenetic tree leading to the New World monkey FPR2. Phylogenetic analysis of positive selection at the codon level in carnivores revealed similar patterns of positive selection in FPR2. The sites under positive selection that appeared in carnivores mapped to the N-terminus and the third and fourth extracellular domain, the exact regions that appear to be undergoing selection in primates as well (Figure 3). To determine how these observed evolutionary changes may influence FPR immune functions, we next took an empirical approach to assess interactions between FPR homologs and known microbial ligands.

**Figure 3.**
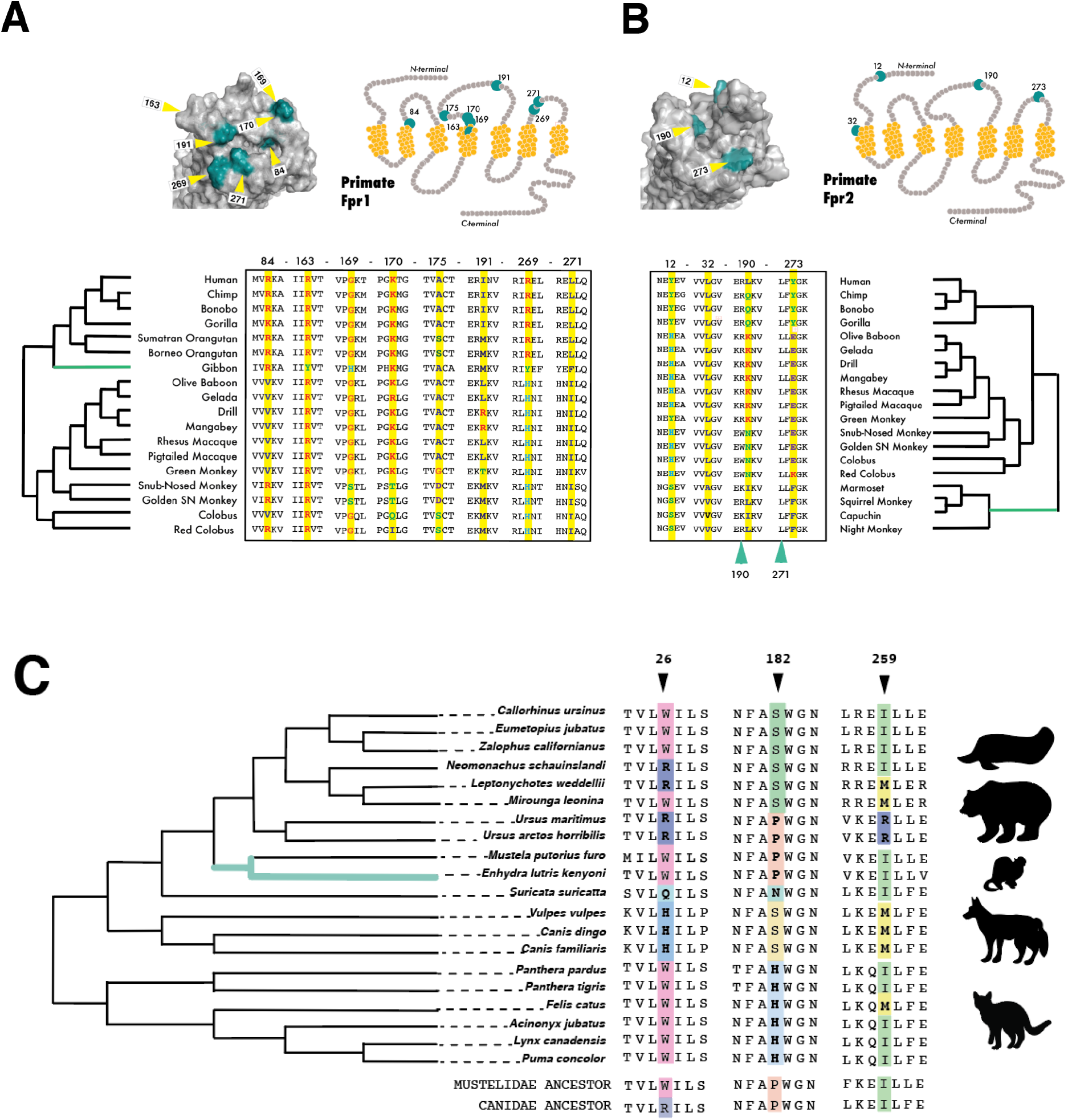
Evidence of repeated positive selection among primate and carnivore FPRs. **A)** Sites in primate FPR1 with elevated dN/dS as determined by PAML and HyPhy. Residues of the transmembrane domain are denoted in yellow on the protein diagram of the FPR1 and FPR2 model, with the majority of high dN/dS sites located in the extracellular ligand-binding loops. **B)** Sites in primate FPR2 with elevated dN/dS as determined by PAML and HyPhy. Residues of the transmembrane domain are denoted in yellow on the protein diagram, with all high dN/dS sites located in the extracellular ligand-binding loops. Branch tests indicate elevated dN/dS in the common ancestor of New World monkeys, which have also lost FPR1. **C)** Sites in carnivore FPR2 with elevated dN/dS as determined by PAML and HyPhy.

### Recognition of bacterial ligands by mammalian FPRs

To assess the functional consequences of sequence variation in primate FPRs, we focused on interactions with the pathogenic bacterium *Staphylococcus aureus* due to its expression of both inhibitors and activators of FPRs (Annette M. Stemerding 20013; Prat 2009; Koymans 2017.; Kretschmer 2010). *S. aureus* is a gram-positive bacterium known for its multiplicity of virulence factors, rapid acquisition of antibiotic resistance, and ability to infect a broad range of mammals beyond humans including livestock, rodents, and companion animals (Thammavongsa 2015; Koymans 2017; Haag et al.). This adaptable microbe colonizes the nares of roughly 30% of humans, but is also a major cause of skin and soft tissue infections, bacterial sepsis, pneumonia, and other life-threatening infections (Thammavongsa 2015.; Haag 2019). *S. aureus* produces many potent toxins that are believed to contribute to its virulence, including enteroxins, leukocidins, and alpha-hemolysin (Balasubramanian 2014.; Lowy 1998; Priatkin and Kuz’menko 2012).

*S. aureus* is able to evade host immune responses in part through the release of proteins that inhibit immune receptors such as TLRs, FPRs, and complement receptors (Thammavongsa 2015.; Prat 2009; Kretschmer 2010; Koymans 2017; Wright 2007) While *S. aureus* is generally believed to be human-adapted, primates, rodents and livestock are frequently colonized by divergent strains of *S. aureus* and related staphylococci (Schaumburg 2012). Many strains sampled from wild gorillas, chimpanzees, green monkeys, and colobus monkeys contain the gene for the virulence factor staphylococcal enterotoxin B (SEB) as well as other virulence factors (Schaumburg 2012). SEB has been shown to potently activate FPRs (Dürr 2006, Kretschmer 2010; Fu 2006). In addition, *S. aureus* produces formylated peptides that can induce cell migration in neutrophils (Dürr 2006). These bacteria also produce a number of molecules that directly bind and inhibit FPRs, such as FLIPr and FLIPr-like which have demonstrated specificity for FPR2, although FLIPr-like has also been shown to bind FPR1 with lower affinity (Stemerding 2013).

We tested binding of proteins fluorescently labeled with fluorescein, FITC-SEB or FITC-FLIPr-like, respectively (Figure 4A,B) to human HEK293T cells expressing primate FPRs by flow cytometry. Binding was assessed based on FITC+ after incubation and washing of unbound labeled protein. Our results for SEB showed evidence of binding for many of the receptors tested. With the exception of human FPR2, all of the primate FPR1 proteins bound to SEB (Figure 4C, D, E, F). The *S. aureus* inhibitor FLIPr-like did not bind FPR proteins with any clear phylogenetic relationship (Figure 4B, C, D, F). Most strikingly, bonobo FPR1 bound FLIPr-like similarly to human FPR2, not human FPR1, despite sharing 98% sequence identity with human FPR1.

**Figure 4.**
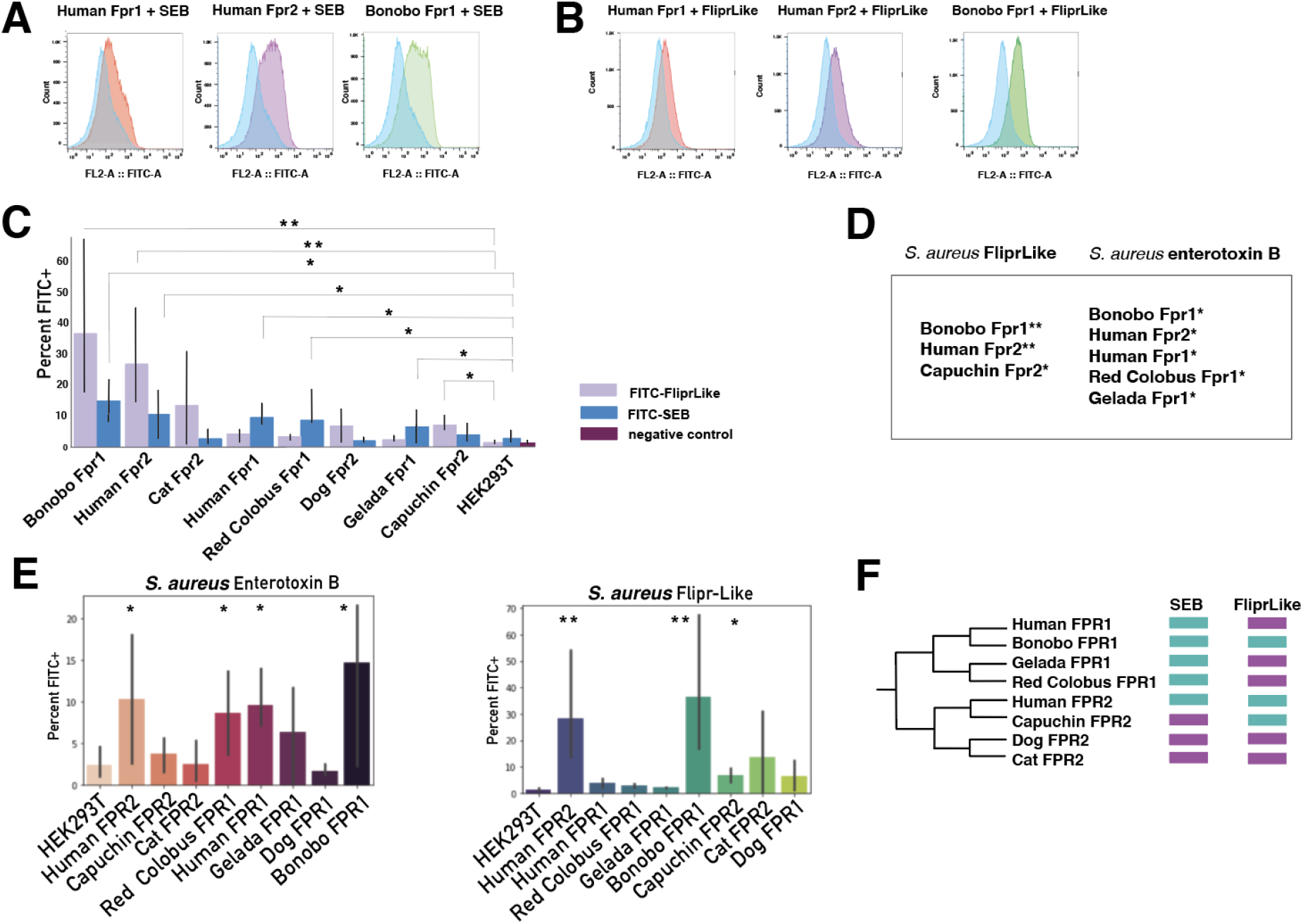
Binding interactions between *S. aureus* ligands and mammalian FPRs. **A)** HEK293T cells expressing mammalian FPRs were incubated with FITC-conjugated *S. aureus* enterotoxin B (SEB), washed, and analyzed by flow cytometry. Data from at least three flow cytometry experiments were collected for each species. Singlet cells were gated and percent of FL2 (488+) positive cells reported from the parent. **B)** HEK293T cells expressing mammalian FPRs were incubated with FITC-conjugated *S. aureus* FLIPr-like, washed, and analyzed by flow cytometry. Data from at least three flow cytometry experiments were collected for each species. Singlet cells were gated and percent of FL2 (488+) positive cells reported from the parent. **C)** Percent FITC+ cells plotted from a subset of singlets, HEK293T cells incubated with FITC-labeled protein used as control to set negative levels. **D)** Table showing FLIPr-like, SEB labeled proteins bound to primate FPRs in this study **E)** Labeled SEB and FLIPr-like proteins bound to primate FPRs, data broken out by ligand **F)** Cladogram of FPRs tested showing positive, statistically significant binding (cyan) or no significant binding (purple)

We next considered which FPR amino acid positions were most likely to be responsible for these binding differences observed. Without crystal structures available for FPRs, we generated homology models using I-TASSER given there are many related GPCRs with sufficient similarity to act as reference structures. These structures were docked using the entire FPR extracellular region with the seven amino acids that form the region of FLIPr and FLIPr-like N-terminal peptide required for its inhibitory activity (Stemerding 2013) using Schrödinger Glide. For FPR1, several amino acids in the first and last extracellular loops formed hydrogen bonds with the FLIPr-like peptide. One of these, site 190, exhibits signatures of positive selection in primates. Moreover, site 190 also differs in both human and red colobus FPR1, neither of which bind to FLIPr-like. Notably, FLIPr-like is also predicted to interact with human FPR2 in the same region important for binding to many other microbial ligands. In fact, there were several sites of overlap between the two including positions S6, N10, E189, and R190 (Figure 5A). Adjacent to this site in human FPR2, site 189 also forms a hydrogen bond with FLIPr-like in docking simulations. There were also several sites of overlap between the two including positions S6, N10, and R190 (Figure 5C). Several studies have shown site 190 can modify peptide binding to other microbial ligands, such as the *S. aureus* chemotaxis inhibitor CHIPs, which shows a preference for lysine over tryptophan at position 190 (Mills 2007). This same site has been shown to modify susceptibility of immune cells to recognition by the *Yersinia pestis* type 3 secretion system, where tryptophan at position 190 has been shown to drastically reduce cell binding versus wildtype K190 (Osei-Owusu 2019).

**Figure 5.**
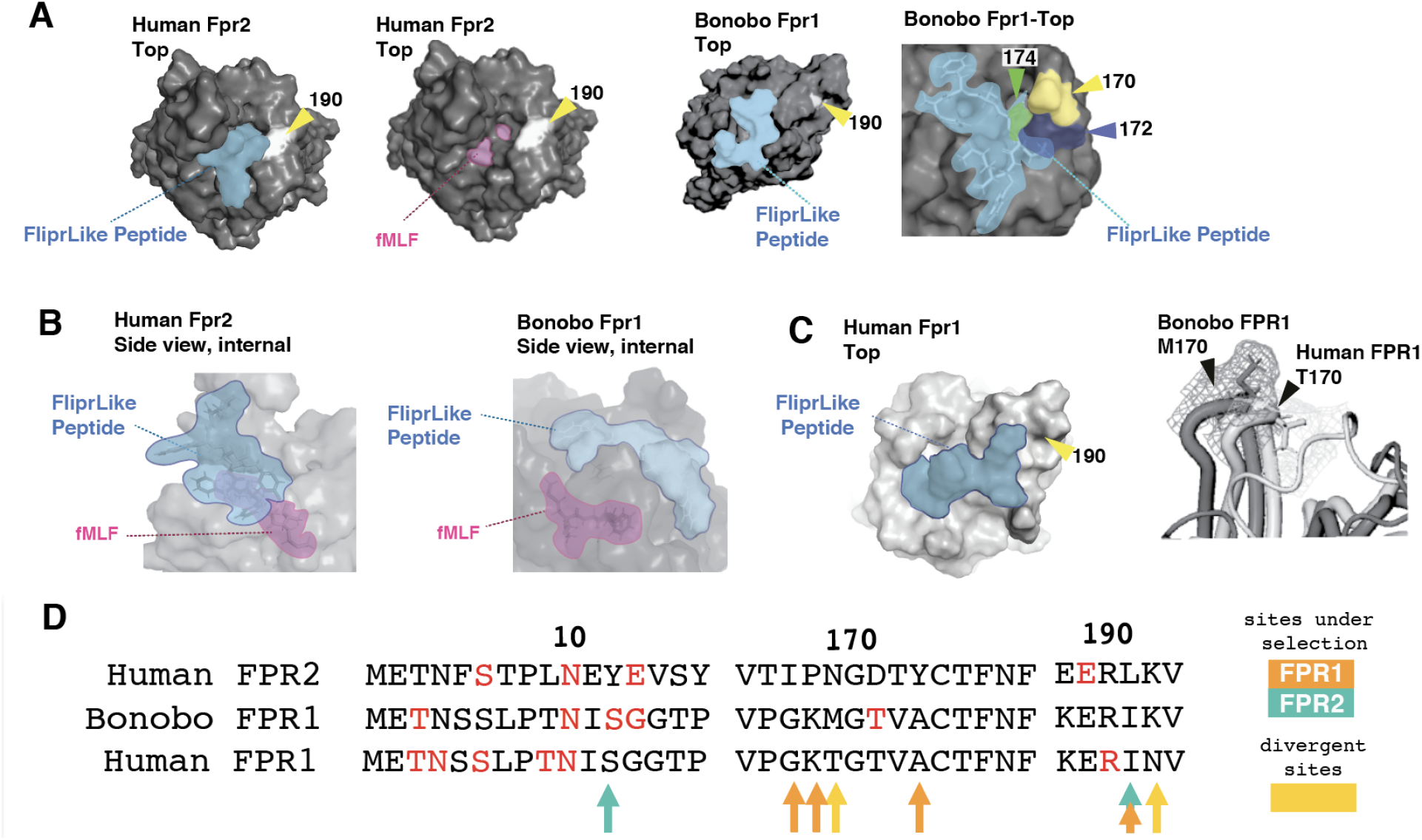
Rapidly-evolving surfaces of FPRs contribute to bacterial ligand binding. **A)** Schrödinger Glide ligand docking prediction reveals that the N-terminal peptide of FLIPr-like, required for inhibitory function, docks adjacent to the canonical FPR ligand fMLF. Site 190 labeled for orientation. **B)** Side view shows the FLIPr-like docking pose internal to the alpha-helices of FPR2, aligned to the fMLF docking pose. **C)** FLIPr-like shows different hydrogen bonding patterns to bonobo FPR1 and docks in a slightly different region of the receptor, due to a protruding ridge formed by M170 in the homology model of bonobo FPR1. Side view of bonobo FPR1 shows similar arrangement between FLIPr-like and fMLF docking, despite different docking pose. **D)** Sites that differ between bonobo FPR1 and human FPR1 are also adjacent to sites predicted to be undergoing positive selection. Sites labeled with red text form hydrogen bonds with FLIPr-like. Arrows in orange indicate sites with high dN/dS for FPR1, green for FPR2. Sites that diverge between bonobo FPR1 and human FPR1 in yellow.

In the bonobo FPR1 docking simulations, the region near sites 189-190 did not interact with the FLIPr-like peptide (Figure 8B). It appears that the amino acid at position 170, which differs between human FPR1 (threonine) and bonobo FPR1 (methionine), creates a ridge where the FLIPr-like peptide docks in the bonobo model. To test how this may affect the inhibitory activity of FLIPr-like, we performed an additional docking simulation with f-MLF, the canonical activator of FPRs. By comparing of the docking poses of human FPR1 and bonobo FPR1, we observed similar binding of the FLIPr-like peptide across several of the extracellular loops of human FPR2 and bonobo FPR1, with the fMLF peptide buried in the central region of the helices (Figure 5A,B). In closing, these results indicate that differences at single rapidly-evolving sites play a large effect in ligand recognition patterns among FPRs, with important consequences for pathogen detection and subsequent infection outcomes.

## DISCUSSION

Human FPR1 can detect formylated peptides at nanomolar concentrations (Mills 2007), which may explain why FPR1 is so highly expressed in human leukocytes (T et al.; Khau 2011; Vacchelli 2020). Since much is known about why FPR genes may expand in specific lineages, it may be worthwhile to seek answers for why genes in this family may undergo loss, such as observed in our study. Loss of FPRs could be beneficial in some contexts, as FPRs have been associated with exacerbating human disease states such as glioblastoma (Yao 2008; Cussell 2009) breast cancer (Khau 2011) and inflammatory diseases ((Cussell 2019, 1; Leslie 2011;; Zhang 2003). Additionally, FPR1 was recently shown to be the receptor for the *Yersinia pestis* type 3 secretion system, a function which can be lost when site 190 is modified from an arginine to a tryptophan (Osei-Owusu 2019). Loss of this gene may confer benefit in animals otherwise susceptible to plague, providing an explanation for the loss of an otherwise highly beneficial gene. Alternatively, redundancy among FPR paralogs within species could lead to gene loss through entirely neutral processes.

Our analysis suggests that FPR1 function was lost early in the New World monkey lineage. Like other immune receptor families including TLRs (Levin and Malik 2017), gene duplication and loss have occurred periodically during the evolution of the FPR gene family in mammals as illustrated by the wide range of copy numbers across taxa. Studies in rodents have revealed remarkable functional plasticity in FPRs (Dietschi 2017). This plasticity likely explains in part why these genes have undergone repeated duplication and loss in so many lineages. There appears to be a broad capacity for FPRs to act as environmental sensors, as demonstrated by aforementioned studies in rodents. In this study we observed several features of a classic molecular arms race dynamic, most notably evidence of repeated positive selection impacting recognition of a microbial pathogen. We pinpointed a rapidly diverging region in the FPRs at sites 190 and 192, which plays a key role in microbial ligand recognition. The *S. aureus* FPR inhibitors CHIPs and FLIPr/FLIPr-like preferentially bind to the FPR1 R190 variant (Mills 2007). It is notable that human FPR2 encodes a charged amino acid glutamate at position 190, possibly reducing its affinity for CHIPs and increasing affinity for FLIPr-like. It is possible that this region is targeted by *S. aureus* inhibitors due to its important role in ligand binding.

Our results indicate that bonobo FPR1 binds to *S. aureus* inhibitor FLIPr-like in a manner similar to human FPR2, even though it shares far more sequence similarity with human FPR1. Notably, this is not the first time that this region has been implicated in interactions with *S. aureus*. We observed several FPR1 single nucleotide polymorphisms in the human population that may influence bacterial ligand recognition. Site R190, for example, is predicted to be involved in binding FLIPr-like. Additionally, the site which forms the ridge in bonobo FPR1, T170M, naturally occurs in the human population, as well as an additional T170P polymorphism which occurs at at low frequency (Figure 6). The consequences of these human FPR variants for recognition by FLIPR-like and other bacterial inhibitors will be an important area for future investigation. Together this work provides a key link between immune receptor variation and recognition by bacterial pathogen virulence factors.

**Figure 6.**
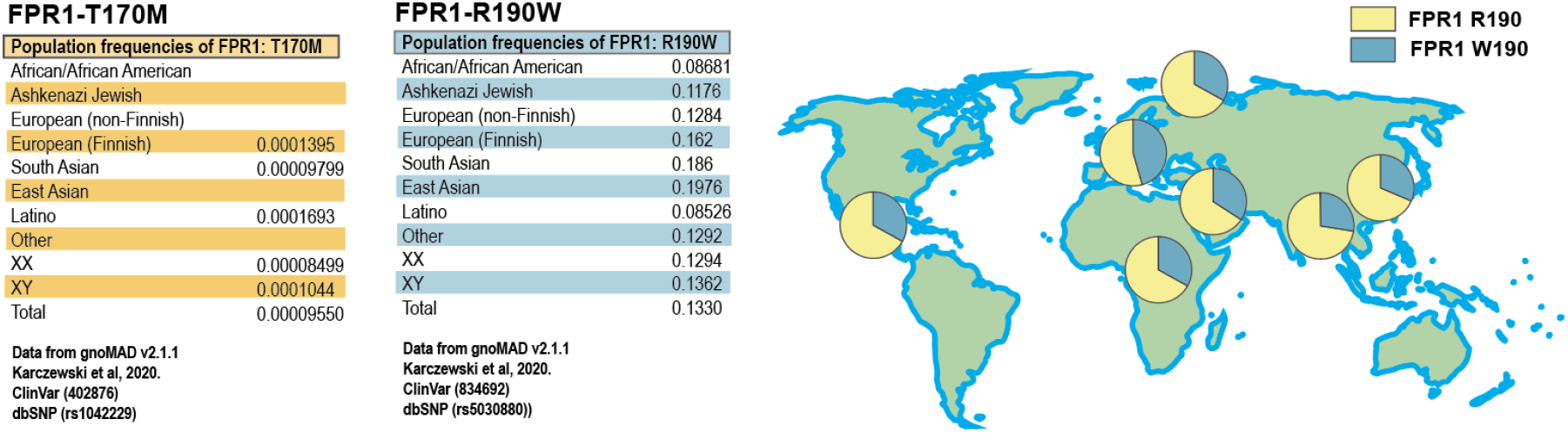
Allele frequencies of abundant human FPR polymorphisms. Data obtained from the gnoMAD server.

## Author Contributions

N.M.P. and M.F.B. conceived the study. N.M.P. performed all phylogenetic analyses, structural modeling, and ligand-binding experiments with assistance from H. A-Z and J.T. FLIPr-like protein was purified by J.R. and supervised by B.V.G. The original manuscript was prepared by N.M.P. with assistance from M.F.B. All authors reviewed and edited the final manuscript.

## Acknowledgements

We are grateful to members of the Barber lab for helpful discussions and feedback on the manuscript. This work was supported by the National Institutes of Health (R35GM133652 to M.F.B., R35GM140852 to B.V.G.). N.M.P. was supported by the National Institutes of Health training grant T32HD07348. The funders played no role in study design, data collection, interpretation, or the decision to publish this study.

## Competing Interests

The authors declare no competing interests.

## METHODS

### Phylogenetic analysis

We inferred amino acid sites exhibiting elevated dN/dS using multiple computational methods. Our dataset included available nucleotide coding sequences (cDNA) of FPR1 for 18 primate species (human, drill, mangabey, red colobus, black and white colobus, snub-nosed monkey, golden snub-nosed monkey, Sumatran orangutan, Bornean orangutan, gorilla, chimpanzee, bonobo, white-cheeked gibbon, green monkey, crab-eating macaque, pig-tailed macaque, gelada, olive baboon), with areas of ambiguity and stop codons removed. A gene tree for FPR paralogs was generated with PhyML (phylogenetics by maximum likelihood) with 1000 bootstraps (Supplemental Figure 1). Potential sites under positive selection were detected using the phylogenetic analysis by maximum likelihood (PAML) package (Yang), which detects signs of positive selection from the frequency of nonsynonymous/synonymous amino acid substitutions at each site (ω = dN/dS) based on maximum likelihood. Additional computational methods MEME (Pond and Frost) and FuBar (Murrell et al.) from the DataMonkey adaptive evolution server were cross-referenced and sites that appeared in more than one analysis with high confidence (p< 0.01) were included. absREL analysis which tests for branch site episodic selection was also performed (Smith et al.).

### Cloning and Lentiviral transduction of FPRs in HEK293T cells

FPR1 genes for human, bonobo, gelada, and red colobus and FPR2 genes for human, capuchin, dog and cat were cloned from cDNA (Human FPR1 and FPR2), synthesized by Genewiz (gelada and red colobus), or synthesized as gBlocks by IDT (capuchin, dog, cat) including Kozak sequence and C-terminal Flag-tag. DNA fragments were subsequently cloned into the pBABE lentiviral vector using SLIC or Gibson cloning methods. Full length FPR1 orthologs were cloned into pBABE vector using the Gibson method for a FLAG epitope tag (Gibson et al., 2009). After expression was verified in cell lines by Western blot using anti Flag tag antibodies (Monoclonal ANTI-FLAG^®^ M1, Sigma Aldrich #F3040), surface expression was verified for FPR1 using Thermo Fisher FPR1 polyclonal antibody ref number PA5-33534) and cell lines with comparable cell surface expression were used for binding experiments. FITC-labeling was performed per manufacturer’s instructions, and the Thermo Scientific^™^ Pierce^™^ Dye Removal Columns, (part number 22858) were used to remove excess dye.

### Flow Cytometry

Cells expressing FPRs from primates (as described above) were counted, suspended at 10^5^ cells/ml i4μg of FITC-labeled FLIPr-like or SEB proteins were incubated in 100μl sterile phosphate buffered saline + 100nM PMSF at 4°C with nutation for 1 hour, washed 3 with 1 ml ice-cold PBS and analyzed on a SONY SH800 flow cytometer and assessed for binding by proportion of cells positive for 488+ as compared to negative control (non-transfected HEK cells incubated with fluorescently labeled proteins in parallel with test samples).

### FLIPr-like Peptide Docking to human FPRs

All structures for analysis were generated using the I-TASSER server (Yang and Zhang), except for f-MLF and FFSYEWK (which represents the first seven amino acids of FLIPr and FLIPr-like proteins) (Annette M. Stemerding et al.) generated in PyMol (Schrodinger). Peptide docking was run with Schrödinger Glide, SP and XP dock with sidechain protonation set to represent charged states at pH 7, highest ranking docking used for analysis.

### Genome scanning for FPR1 pseudogenes in New World Monkeys

Complete genomes for available New World Monkey species were queried for FPR1 pseudogenes by BLAST search, BLAT search of genome in UCSC Genome Browser (*Callithrix jacchus*, *Saimiri boliviensis*) while pseudogenes for *Cebus capucinus imitator*, *Sapajus apella*, and *Aotus nancymaae* were identified by BLAT search for similarity in genome viewer in NCBI’s Genome Data viewer. Details for genome assemblies in Supplemental Figure 2. Pseudogenes were aligned using MUSCLE. Resulting data was analyzed by alignment and exons were searched for and in some cases eliminated using GENESCAN (Burge and Karlin 1997).

## Notes

### Competing Interest Statement

The authors have declared no competing interest.

## REFERENCES

Aleru, Omoshola, and Matthew F. Barber. “Battlefronts of Evolutionary Conflict between Bacteria and Animal Hosts.” PLoS Pathogens, vol. 16, no. 9, Sept. 2020. PubMed Central, doi:10.1371/journal.ppat.1008797.

Balasubramanian, Divya, Lamia Harper, Bo Shopsin, and Victor J. Torres. 2017. “Staphylococcus Aureus Pathogenesis in Diverse Host Environments.” Pathogens and Disease 75 (1). https://doi.org/10.1093/femspd/ftx005.

Bao, L., et al. “Mapping of Genes for the Human C5a Receptor (C5AR), Human FMLP Receptor (FPR), and Two FMLP Receptor Homologue Orphan Receptors (FPRH1, FPRH2) to Chromosome 19.” Genomics, vol. 13, no. 2, June 1992, pp. 437–40. PubMed, doi:10.1016/0888-7543(92)90265-t.

Barber, Matthew F., and Nels C. Elde. “Escape from Bacterial Iron Piracy through Rapid Evolution of Transferrin.” Science (New York, N.Y.), vol. 346, no. 6215, Dec. 2014, pp. 1362–66. PubMed Central, doi:10.1126/science.1259329.

Brunette, Rebecca L., et al. “Extensive Evolutionary and Functional Diversity among Mammalian AIM2-like Receptors.” The Journal of Experimental Medicine, vol. 209, no. 11, Oct. 2012, pp. 1969–83. PubMed Central, doi:10.1084/jem.20121960.

Bufe, Bernd, et al. “Recognition of Bacterial Signal Peptides by Mammalian Formyl Peptide Receptors: A NEW MECHANISM FOR SENSING PATHOGENS*.” Journal of Biological Chemistry, vol. 290, no. 12, Mar. 2015, pp. 7369–87. ScienceDirect, doi:10.1074/jbc.M114.626747.

Burge, C., and S. Karlin. “Prediction of Complete Gene Structures in Human Genomic DNA.” Journal of Molecular Biology, vol. 268, no. 1, Apr. 1997, pp. 78–94. PubMed, doi:10.1006/jmbi.1997.0951.

Crouser, Elliott D., et al. “Monocyte Activation by Necrotic Cells Is Promoted by Mitochondrial Proteins and Formyl Peptide Receptors.” Critical Care Medicine, vol. 37, no. 6, June 2009, pp. 2000–09. PubMed, doi:10.1097/CCM.0b013e3181a001ae.

Cussell, Peter J. G., et al. “The Formyl Peptide Receptor Agonist FPRa14 Induces Differentiation of Neuro2a Mouse Neuroblastoma Cells into Multiple Distinct Morphologies Which Can Be Specifically Inhibited with FPR Antagonists and FPR Knockdown Using SiRNA.” PLoS ONE, vol. 14, no. 6, Public Library of Science, June 2019, pp. e0217815–e0217815. go-gale-com.libproxy.uoregon.edu, doi:10.1371/journal.pone.0217815.

Daugherty, Matthew D., and Harmit S. Malik. “Rules of Engagement: Molecular Insights from Host-Virus Arms Races.” Annual Review of Genetics, vol. 46, 2012, pp. 677–700. PubMed, doi:10.1146/annurev-genet-110711-155522.

Dietschi, Quentin, et al. “Evolution of Immune Chemoreceptors into Sensors of the Outside World.” Proceedings of the National Academy of Sciences, National Academy of Sciences, June 2017. www.pnas.org, doi:10.1073/pnas.1704009114.

Dürr, Manuela C., et al. “Neutrophil Chemotaxis by Pathogen-Associated Molecular Patterns--Formylated Peptides Are Crucial but Not the Sole Neutrophil Attractants Produced by Staphylococcus Aureus.” Cellular Microbiology, vol. 8, no. 2, Feb. 2006, pp. 207–17. PubMed, doi:10.1111/j.1462-5822.2005.00610.x.

Fu, Huamei, et al. “Ligand Recognition and Activation of Formyl Peptide Receptors in Neutrophils.” Journal of Leukocyte Biology, vol. 79, no. 2, 2006, pp. 247–56. Wiley Online Library, doi:https://doi.org/10.1189/jlb.0905498.

Haag, Andreas F., J. Ross Fitzgerald, and José R. Penadés. 2019. “Staphylococcus Aureus in Animals.” Microbiology Spectrum 7 (3).

Harris, R. Alan, et al. “Unusual Sequence Characteristics of Human Chromosome 19 Are Conserved across 11 Nonhuman Primates.” BMC Evolutionary Biology, vol. 20, no. 1, 1, BioMed Central, Dec. 2020, pp. 1–12. link-springer-com.libproxy.uoregon.edu, doi:10.1186/s12862-020-1595-9.

Hughes, Austin L., and Helen Piontkivska. “Functional Diversification of the Toll-like Receptor Gene Family.” Immunogenetics, vol. 60, no. 5, May 2008, pp. 249–56. PubMed Central, doi:10.1007/s00251-008-0283-5.

Hughes, Austin, and Masatoshi Nei. “Hughes AL, Nei M. Evolutionary Relationships of Class II MHC Genes in Mammals. Mol Biol Evol 7: 491.” Molecular Biology and Evolution, vol. 7, Dec. 1990, pp. 491–514.

John, C. D., et al. “Formyl Peptide Receptors and the Regulation of ACTH Secretion: Targets for Annexin A1, Lipoxins, and Bacterial Peptides.” The FASEB Journal: Official Publication of the Federation of American Societies for Experimental Biology, vol. 21, no. 4, Apr. 2007, pp. 1037–46. PubMed Central, doi:10.1096/fj.06-7299com.

Karlsson, Jennie, et al. “The FPR2-Specific Ligand MMK-1 Activates the Neutrophil NADPH-Oxidase, but Triggers No Unique Pathway for Opening of Plasma Membrane Calcium Channels.” Cell Calcium, vol. 45, no. 5, May 2009, pp. 431–38. DOI.org (Crossref), doi:10.1016/j.ceca.2009.02.002.

Karolchik, Donna, et al. “The UCSC Genome Browser.” Current Protocols in Bioinformatics / Editoral Board, Andreas D. Baxevanis… [et Al.], vol. CHAPTER, Dec. 2009, p. Unit1.4. PubMed Central, doi:10.1002/0471250953.bi0104s28.

Khau, Thippadey, et al. “Annexin-1 Signals Mitogen-stimulated Breast Tumor Cell Proliferation by Activation of the Formyl Peptide Receptors (FPRs) 1 and 2.” The FASEB Journal, vol. 25, no. 2, WILEY, 2011, pp. 483–96. alliance-primo.com, doi:10.1096/fj.09-154096.

Kim, Sang Doo, et al. “Functional Expression of Formyl Peptide Receptor Family in Human NK Cells.” The Journal of Immunology, vol. 183, no. 9, American Association of Immunologists, Nov. 2009, pp. 5511–17. www-jimmunol-org.libproxy.uoregon.edu, doi:10.4049/jimmunol.0802986.

Koymans, Kirsten J., et al. “Staphylococcal Immune Evasion Proteins: Structure, Function, and Host Adaptation.” Staphylococcus Aureus: Microbiology, Pathology, Immunology, Therapy and Prophylaxis, edited by Fabio Bagnoli et al., Springer International Publishing, 2017, pp. 441–89. Springer Link, doi:10.1007/82_2015_5017.

Kretschmer, Dorothee, et al. “Human Formyl Peptide Receptor 2 (FPR2/ALX) Senses Highly Pathogenic Staphylococcus Aureus.” Cell Host & Microbe, vol. 7, no. 6, June 2010, pp. 463–73. PubMed Central, doi:10.1016/j.chom.2010.05.012.

Le, Yingying, et al. “Formyl-Peptide Receptors Revisited.” Trends in Immunology, vol. 23, no. 11, Nov. 2002, pp. 541–48. www.cell.com, doi:10.1016/S1471-4906(02)02316-5.

Leslie, Jack, et al. “FPR-1 Is an Important Regulator of Neutrophil Recruitment and a Tissue-Specific Driver of Pulmonary Fibrosis.” JCI Insight, vol. 5, no. 4. PubMed Central, doi:10.1172/jci.insight.125937. Accessed 14 Mar. 2021.

Levin, Tera C., and Harmit S. Malik. “Rapidly Evolving Toll-3/4 Genes Encode Male-Specific Toll-Like Receptors in Drosophila.” Molecular Biology and Evolution, vol. 34, no. 9, Sept. 2017, pp. 2307–23. PubMed, doi:10.1093/molbev/msx168.

Lowy, F. D. 1998. “Staphylococcus Aureus Infections.” The New England Journal of Medicine 339 (8): 520–32. https://doi.org/10.1056/NEJM199808203390806.

McLaughlin, Richard N., and Harmit S. Malik. “Genetic Conflicts: The Usual Suspects and Beyond.” The Journal of Experimental Biology, vol. 220, no. 1, Jan. 2017, pp. 6–17. PubMed Central, doi:10.1242/jeb.148148.

Mills, John S. 2007. “Differential Activation of Polymorphisms of the Formyl Peptide Receptor by Formyl Peptides.” Biochimica Et Biophysica Acta 1772 (9): 1085–92. https://doi.org/10.1016/j.bbadis.2007.06.001.

Muto, Yoshinori, et al. “Adaptive Evolution of Formyl Peptide Receptors in Mammals.” Journal of Molecular Evolution, vol. 80, no. 2, Feb. 2015, pp. 130–41. PubMed, doi:10.1007/s00239-015-9666-z.

Önnheim, Karin, et al. “A Novel Receptor Cross-Talk between the ATP Receptor P2Y2 and Formyl Peptide Receptors Reactivates Desensitized Neutrophils to Produce Superoxide.” Experimental Cell Research, vol. 323, no. 1, Apr. 2014, pp. 209–17. ScienceDirect, doi:10.1016/j.yexcr.2014.01.023.

Osei-Owusu, Patrick, et al. “FPR1 Is the Plague Receptor on Host Immune Cells.” Nature, vol. 574, no. 7776, 2019, pp. 57–62. PubMed, doi:10.1038/s41586-019-1570-z.

Paterson, Nicole M., et al. “Diversification of CD1 Molecules Shapes Lipid Antigen Selectivity.” Molecular Biology and Evolution, no. msab022, Feb. 2021. Silverchair, doi:10.1093/molbev/msab022.

Pike, MC, et al. “Development of Specific Receptors for N-Formylated Chemotactic Peptides in a Human Monocyte Cell Line Stimulated with Lymphokines.” The Journal of Experimental Medicine, vol. 152, no. 1, July 1980, pp. 31–40.

Prat, Cristina, et al. “A Homolog of Formyl Peptide Receptor-like 1 (FPRL1) Inhibitor from Staphylococcus Aureus (FPRL1 Inhibitory Protein) That Inhibits FPRL1 and FPR.” Journal of Immunology (Baltimore, Md.: 1950), vol. 183, no. 10, Nov. 2009, pp. 6569–78. PubMed, doi:10.4049/jimmunol.0801523.

Priatkin, R. G., and O. M. Kuz’menko. 2010. “[Secreted proteins of Staphylococcus aureus].” Zhurnal Mikrobiologii, Epidemiologii I Immunobiologii, no. 4 (August): 118–24.

Sawyer, Sara L., et al. Positive Selection of Primate TRIM5␣ Identifies a Critical Species-Specific Retroviral Restriction Domain. p. 6. 2832–2837 PNAS. February 22, 2005 vol. 102 no. 8

Schaumburg, F., A. S. Alabi, R. Köck, A. Mellmann, P. G. Kremsner, C. Boesch, K. Becker, F. H. Leendertz, and G. Peters. 2012. “Highly Divergent Staphylococcus Aureus Isolates from African Non-Human Primates.” Environmental Microbiology Reports 4 (1): 141–46. https://doi.org/10.1111/j.1758-2229.2011.00316.x.

Schepetkin, I. A., et al. “Development of Small Molecule Non-Peptide Formyl Peptide Receptor (FPR) Ligands and Molecular Modeling of Their Recognition.” Current Medicinal Chemistry, vol. 21, no. 13, 2014, pp. 1478–504. PubMed Central, doi:10.2174/0929867321666131218095521.

Schiffmann, E., et al. “N-Formylmethionyl Peptides as Chemoattractants for Leucocytes.” Proceedings of the National Academy of Sciences of the United States of America, vol. 72, no. 3, Mar. 1975, pp. 1059–62.

Shimizu, Nobuaki, et al. “A Formylpeptide Receptor, FPRL1, Acts as an Efficient Coreceptor for Primary Isolates of Human Immunodeficiency Virus.” Retrovirology, vol. 5, no. 1, June 2008, p. 52. BioMed Central, doi:10.1186/1742-4690-5-52.

Stemerding, Annette, et al. “Staphylococcus Aureus Formyl Peptide Receptor–like 1 Inhibitor (FLIPr) and Its Homologue FLIPr-like Are Potent FcγR Antagonists That Inhibit IgG-Mediated Effector Functions.” The Journal of Immunology, vol. 191, no. 1, American Association of Immunologists, July 2013, pp. 353–62. www-jimmunol-org.libproxy.uoregon.edu, doi:10.4049/jimmunol.1203243.

Thammavongsa, Vilasack, Hwan Keun Kim, Dominique Missiakas, and Olaf Schneewind. 2015. “Staphylococcal Manipulation of Host Immune Responses.” Nature Reviews. Microbiology 13 (9): 529–43. https://doi.org/10.1038/nrmicro3521.

Thierry-Mieg, Danielle, and Jean Thierry-Mieg. “AceView: A Comprehensive CDNA-Supported Gene and Transcripts Annotation.” Genome Biology, vol. 7, no. 1, Aug. 2006, p. S12. BioMed Central, doi:10.1186/gb-2006-7-s1-s12.

Vacchelli, Erika, Julie Le Naour, and Guido Kroemer. 2020. “The Ambiguous Role of FPR1 in Immunity and Inflammation.” Oncoimmunology 9 (1): 1760061. https://doi.org/10.1080/2162402X.2020.1760061.

van der Lee, Robin, et al. “Genome-Scale Detection of Positive Selection in Nine Primates Predicts Human-Virus Evolutionary Conflicts.” Nucleic Acids Research, vol. 45, no. 18, Oct. 2017, pp. 10634–48. PubMed, doi:10.1093/nar/gkx704.

Wright, Andrew J., Adrian Higginbottom, Didier Philippe, Abhishek Upadhyay, Stefan Bagby, Robert C. Read, Peter N. Monk, and Lynda J. Partridge. 2007. “Characterisation of Receptor Binding by the Chemotaxis Inhibitory Protein of Staphylococcus Aureus and the Effects of the Host Immune Response.” Molecular Immunology 44 (10–4): 2507–17. https://doi.org/10.1016/j.molimm.2006.12.022.

Yang, Huan, et al. “Relating Gene Expression Evolution with CpG Content Changes.” BMC Genomics, vol. 15, no. 1, Aug. 2014, p. 693. BioMed Central, doi:10.1186/1471-2164-15-693.

Yang, Hui, and Peng Shi. “Molecular and Evolutionary Analyses of Formyl Peptide Receptors Suggest the Absence of VNO-Specific FPRs in Primates.” Journal of Genetics and Genomics, vol. 37, no. 12, Dec. 2010, pp. 771–78. DOI.org (Crossref), doi:10.1016/S1673-8527(09)60094-1.

Yang, Z. “PAML 4: Phylogenetic Analysis by Maximum Likelihood.” Molecular Biology and Evolution, vol. 24, no. 8, Apr. 2007, pp. 1586–91. DOI.org (Crossref), doi:10.1093/molbev/msm088.

Yao, Xiao-Hong, et al. “Production of Angiogenic Factors by Human Glioblastoma Cells Following Activation of the G-Protein Coupled Formyl peptide Receptor FPR.” Journal of Neuro-Oncology, vol. 86, no. 1, Jan. 2008, pp. 47–53. Springer Link, doi:10.1007/s11060-007-9443-y.

Zigmond, S H. 1981. “Consequences of Chemotactic Peptide Receptor Modulation for Leukocyte Orientation.” Journal of Cell Biology 88 (3): 644–47. https://doi.org/10.1083/jcb.88.3.644.

